# Competitive asymmetry confers polyploid advantage under environmental stress

**DOI:** 10.1101/2021.11.08.467667

**Authors:** Wen Guo, Na Wei, Guang-You Hao, Shi-Jian Yang, Zhi-Yong Zhu, Yong-Ping Yang, Yuan-Wen Duan

## Abstract

- Competitive asymmetry across heterogeneous environments is crucial for the success of polyploid plants, however, little is known about it. As the major force in plant evolution, polyploids are predicted to maintain the competitive dominance relative to diploids under increased stress conditions.
- To evaluate the hypothesis of competitive asymmetry, we competed tetraploid and diploid plants of perennial herbaceous *Chrysanthemum indicum* L. (Asteraceae) at different relative frequencies under low and high water stress. We quantified the interaction intensity between competing plants of the same (intraploidy) and different ploidy levels (interploidy), and measured functional traits related to gas exchange and plant water use to understand the underlying mechanisms.
- Stronger competitive effects of the tetraploid on diploid provided evidence for the competitive asymmetry between polyploid and diploid plants in *C. indicum*. Such competitive asymmetry was not only maintained under drought (increased water stress), but also translated into higher fitness of the tetraploid consistently across water stress conditions. Functional traits associated with fast growth and efficient water use likely explained the competitive dominance of the tetraploid.
- These results will advance our understanding of species interactions between polyploid and diploid plants, and provide insights into population dynamics and species distribution under environmental change.

## Introduction

Competitive interaction and habitat filtering (e.g. imposed by stress) influence species niche and distribution and community structure (Seabloom *et al*., 2003; Maire *et al*., 2012; Napier *et al*., 2016). These effects can be pronounced especially during incipient speciation, where the newly evolved taxon is at a numeric disadvantage when competing with abundant, locally adapted parental taxa for the same essential resources both abiotically and biotically (Levin, 1975; Fowler & Levin, 1984; Schluter, 2001; Fowler & Levin, 2016). Such minority disadvantage is thought to be common during polyploid speciation (Levin, 1975) and can be prevalent in flowering plants, as polyploidy (or whole-genome duplication) plays a critical role in speciation events of angiosperms (Wood *et al*., 2009) and has a widespread incidence across numerous plant lineages (Jiao *et al*., 2011). To overcome the minority effects, increased niche differentiation and competitive dominance over diploid progenitors are predicted to be important for polyploid establishment and evolution (Fowler & Levin, 1984, 2016). Both conditions could lead to niche and distribution divergence (e.g. allopatry) between polyploids and diploids or fine-scale spatial segregation in the cases of niche conservatism and sympatry, as seen across diverse plant lineages (Glennon *et al*., 2014; Marchant *et al*., 2016; Wei *et al*., 2017; López-Jurado *et al*., 2019). As a result of independent evolution in allopatry or spatial segregation in sympatry, it remains largely unknown whether contemporary polyploids are competitively dominant showing fitness advantage across environments (Wei *et al*., 2019) when competing with diploids that have evolved without polyploid competitors (Collins *et al*., 2011; Thompson *et al*., 2015; Rey *et al*., 2017). This question is of critical relevance to predicting species interactions and range shifts under changing climates, where tracking moving climatic envelopes likely increase species encounter between polyploid and diploid taxa.

Competitive interactions are predicted to be asymmetric between polyploids and diploids, with polyploids being competitively superior owing to enhanced genomic diversity and versatility (Otto & Whitton, 2000; Ramsey & Ramsey, 2014; Soltis *et al*., 2016; Van de Peer *et al*., 2017; Wei *et al*., 2019, 2020). Thus, the competitive effect of polyploids on diploids (α_2*x*,4*x*_, *4x* and 2*x* refer to tetraploids and diploids as an example hereafter; Fig. 1a,b, blue solid line and blue square) is expected to be stronger than the effect of diploids on polyploids (α_4*x*,2*x*_; Fig. 1a,b, red solid line and red circle). Likewise, if polyploids are better competitors than diploids, polyploids are expected to experience a higher reduction in fitness when competing with individuals of its own (intraploidy) than they do with individuals of diploids (interploidy). In other words, polyploids should be limited more by intraploidy than interploidy competition (α_4*x*,4*x*_ > α_4*x*,2*x*_; Fig. 1c). In contrast, competitively inferior diploids are expected to be limited more by interploidy than intraploidy competition (α_2*x*,4*x*_ > α_2*x*,2*x*_; Fig. 1c). While quantifying intra- vs. interploidy competition is fundamental to understanding the magnitude and outcome of species interactions, few experimental studies have done so between polyploid and diploid competitors (Collins *et al*., 2011; Thompson *et al*., 2015; Rey *et al*., 2017).

**Fig. 1.**
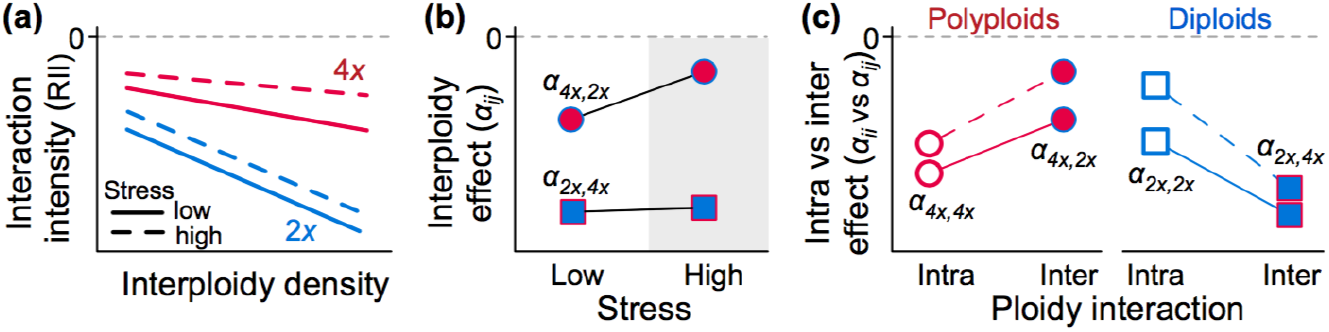
Hypotheses of competitive asymmetry between polyploids and diploids. (a) The competitive effect (i.e. slope) and competitive response (*y*-axis) of interploidy density are expected to be lower on polyploids (e.g. 4*x*, tetraploids) relative to diploids (2*x*), if the 4*x* is a stronger competitor than the 2*x*. Plant-plant interaction is quatified using the relative interaction index (−1 ≤ RII ≤ 1), with RII < 0 indicating competition. (b) The interploidy competitive effect represents the slope (α*_ij_*, where *i* and *j* refer to the 2*x* or 4*x*, respectively) in (a). Stress is expected to lower competitive effect, especially the competitive effect of 2*x* on 4*x* (α_4*x*,2*x*_, circles), but less so for the competitive effect of 4*x* on 2*x* (α_2*x*,4*x*_, squares), if the 4*x* is competitively superior and less affected by stress relative to the 2*x*. As a result, the competitive asymmetry can be maintained or even enhanced under stress. (c) If the 4*x* is a stronger competitor than the 2*x*, the 4*x* is expected to be limited more by intraploidy effect (α_4*x*,4*x*_) than interploidy effect (α_4*x*,2*x*_). This pattern is likely more pronouced under stress, if the competitively inferior 2*x* is affected more by stress than the 4*x*, leading to stronger reduction in α_4*x*,2*x*_ (filled red circles) than α_4*x*,4*x*_ (open red circles). The opposite pattern is expected for the 2*x* (blue squares).

Competitive interactions can be environment dependent (Brooker, 2006; Ploughe *et al*., 2019; Zhang & Tielböerger, 2020). The stress gradient hypothesis (SGH; Bertness & Callaway, 1994; Maestre *et al*., 2009) provides a framework for predicting how the magnitude of intraploidy and interploidy competitions change under environmental stress. The SGH predicts that competitive interactions weaken as the environment becomes more stressful, because individual competitors that are under stress likely affect each other less than under low stress conditions. Thus, the interaction intensity response curve (Fig. 1a) is expected to move upwards with increased stress levels (Zhang & Tielböerger, 2020). Yet, the extent to which stress lowers competitive effects is expected to differ between polyploids and diploids (Fig. 1a,b), due to their differential abilities of stress tolerance. Polyploids that are often more stress tolerant than diploids (Van de Peer *et al*., 2021) are likely impacted less by stress, and thus their competitive effect on diploids (α_2*x*,4*x*_) is expected to be reduced less than the effect of diploids on polyploids (α_4*x*,2*x*_; Fig. 1b). As a result, competitive asymmetry between polyploids and diploids is likely maintained or enhanced under stress (Fig. 1b); this hypothesis is yet to be tested.

Functional trait divergence can influence the magnitude of competitive interactions (Gaudet & Keddy, 1988; Kraft *et al*., 2015) and thus is expected to underlie the hypothesized competitive asymmetry between polyploids and diploids. Maintaining the asymmetry across environments, nevertheless, depends on the extent to which traits respond adaptively to changing environments (Wei *et al*., 2019). Under favorable, low-stress conditions, competition favors an acquisitive strategy that can enhance resource consumption and carbon assimilation (Wei *et al*., 2019; Lorts & Lasky, 2020), whereas under stressful conditions more conservative strategy is needed (Wei *et al*., 2019). For instance, polyploids can exhibit higher gas exchanges (net photosynthesis, stomatal conductance, transpiration rate) that can promote plant growth than diploids under low drought stress (Vyas *et al*., 2007). Under high drought stress, polyploids can be more drought tolerant than diploids, due to, for instance, higher water use efficiency (Manzaneda *et al*., 2012) and xylem structure that permits water transport and photosynthesis under very negative water potential (Li *et al*., 1996; Hao *et al*., 2013). Thus, linking functional traits to competitive ability across environments is critical for a mechanistic understanding of competitive interactions between polyploids and diploids.

To evaluate the hypothesis of competitive asymmetry between polyploids and diploids under changing environments and the underlying mechanisms, we competed the diploid and tetraploid plants of a perennial herbaceous species complex, *Chrysanthemum indicum* L. (Asteraceae), at different relative frequencies under low and high water stress. The diploid and tetraploid populations of *C. indicum* in China are mainly in allopatry, with the tetraploids that likely experienced a geological period of drought during the Quaternary glaciation (Yang *et al*., 2006; Li *et al*., 2014) more widespread. In their broad sympatric portion in central China, the diploid and tetraploid populations are spatially separated. Here, by quantifying intraploidy and interploidy competitive effects and a suite of functional traits related to gas exchange and plant water use, we aimed to address three core questions: (1) is there competitive asymmetry between polyploids and diploids with polyploids being competitively dominant? If so, (2) can competitive asymmetry be maintained or even enhanced under water stress (drought)? and (3) how do functional traits explain the competitive dominance of polyploids over diploids?

## Materials and Methods

### Study system

*Chrysanthemum indicum* grows primarily in open and xeric habitats. This species complex comprises two main ploidy levels, diploid (2*n* = 18 chromosomes) and tetraploid (2*n* = 36) that likely has both autopolyploid and allopolyploid origins (Yang *et al*., 2006). Within their sympatric distribution in Shennongjia, China (31.74472°N, 110.67583°E), we collected seeds from four 2*x* populations and six 4*x* populations (Table S1) in November 2015. Specifically, 5–10 flower heads were collected from individual maternal plants each population (*N* = 3–5 or 6–10 plants for the 2*x* and 4*x* populations, respectively, due to variation in population sizes). In addition, three fresh leaves were collected from each maternal plant in moist plastic bags for ploidy confirmation at the Kunming Institute of Botany, using flow cytometry following the previous protocol (Guo *et al*., 2016).

### Seedling cultivation

In March 2016, seeds of the 2*x* and 4*x* maternal plants were sowed in peat substrates (0– 10 mm; Novarbo, Lauttakylantie, Finland) in 8 cm pots (dimeter: 8 cm [top] and 6 cm [bottom]; height: 8 cm). Each pot contained *c*. 10 seeds with a total of 60 pots for the 2*x* and 72 pots for the 4*x*. Seedlings were grown under 25:15°C day : night temperatures in a naturally lit glasshouse at the Germplasm Bank of Wild Species (Kunming Institute of Botany, China). After one month, we harvested leaves from three seedlings each pot to confirm ploidy levels. Pots with mixed ploidy levels (*N* = 8 pots) were excluded from this study. In May 2016, seedlings of similar size (*c*. 5 cm in height) were used for the competition experiment and functional traits experiment at the same time.

### Competition experiment

To examine the competitive effects of intraploidy and interploidy density, the 2*x* and 4*x* seedlings were grown together at different relative frequencies (varying in 0, 2, 3 or 4 plants per ploidy level each pot; Fig. S1): 0:2, 0:3, 0:4, 2:0, 2:2, 2:3, 2:4, 3:0, 3:2, 3:3, 3:4, 4:0, 4:2, 4:3, 4:4. These 15 density combinations constituted the basic unit of the competition experiment (Fig. S1) and were replicated 10 times, with 720 total plants in 150 7-L pots filled with 1.5 kg custom potting mixture (1:1, peat : perlite). Water content was maintained at the field capacity (‘FC’; 0.9 kg water per 1.5 kg potting mixture) by watering every two days for 3 wk. On June 1, 2016, we started the water stress treatments, with 5 replicates receiving low water stress treatment (80% FC) and 5 replicates receiving high water stress treatment (20% FC) for 2 months until plant harvesting. To maintain the specific water content, pots were weighted every two days to record water loss and re-watered accordingly. This experiment used a split plot design for the ease of watering. Pots in the glasshouse were not rotated during the experiment due to large pot sizes, and thus pot positions were considered in data analyses to account for potential influence of microenvironment variation. On July 30, 2016, we harvested the aboveground plant materials by ploidy level each pot, but not the belowground due to challenges in separating roots between the 2*x* and 4*x* plants. Trials with the 2*x* and 4*x* plants growing separately indicated similar root to shoot ratios, and thus aboveground dry biomass is expected to correlate closely with whole plant dry biomass in *C. indicum*. The aboveground plant materials were dried at 80°C for 72 h to obtain the average aboveground dry biomass of the 2*x* and 4*x* each pot. Due to occasional plant mortality during the competition experiment, pots (*N* = 10) that did not match their original density combination were excluded from data analyses.

### Functional traits experiment

We measured a suite of functional traits that are relevant to plant growth and drought tolerance (Table 1) in a separate experiment, to avoid potential influence of plant tissue collection on the competition experiment. In the functional traits experiment, 112 seedlings of each ploidy level were grown in 56 pots (2 seedlings of the same ploidy per pot), with 112 total pots for the 2*x* and 4*x*. Ten ‘empty’ pots filled with the same potting mixture but no seedlings were included for measuring evaporation. As in the competition experiment, on June 1, 2016, half of the experimental (*N* = 28 pots per ploidy level) and ‘empty’ pots (*N* = 5) were assigned at random to the low water stress treatment (80% FC) and half to the high water stress treatment (20% FC) for 2 months. No seedling mortality occurred during this experiment.

**Table 1.**
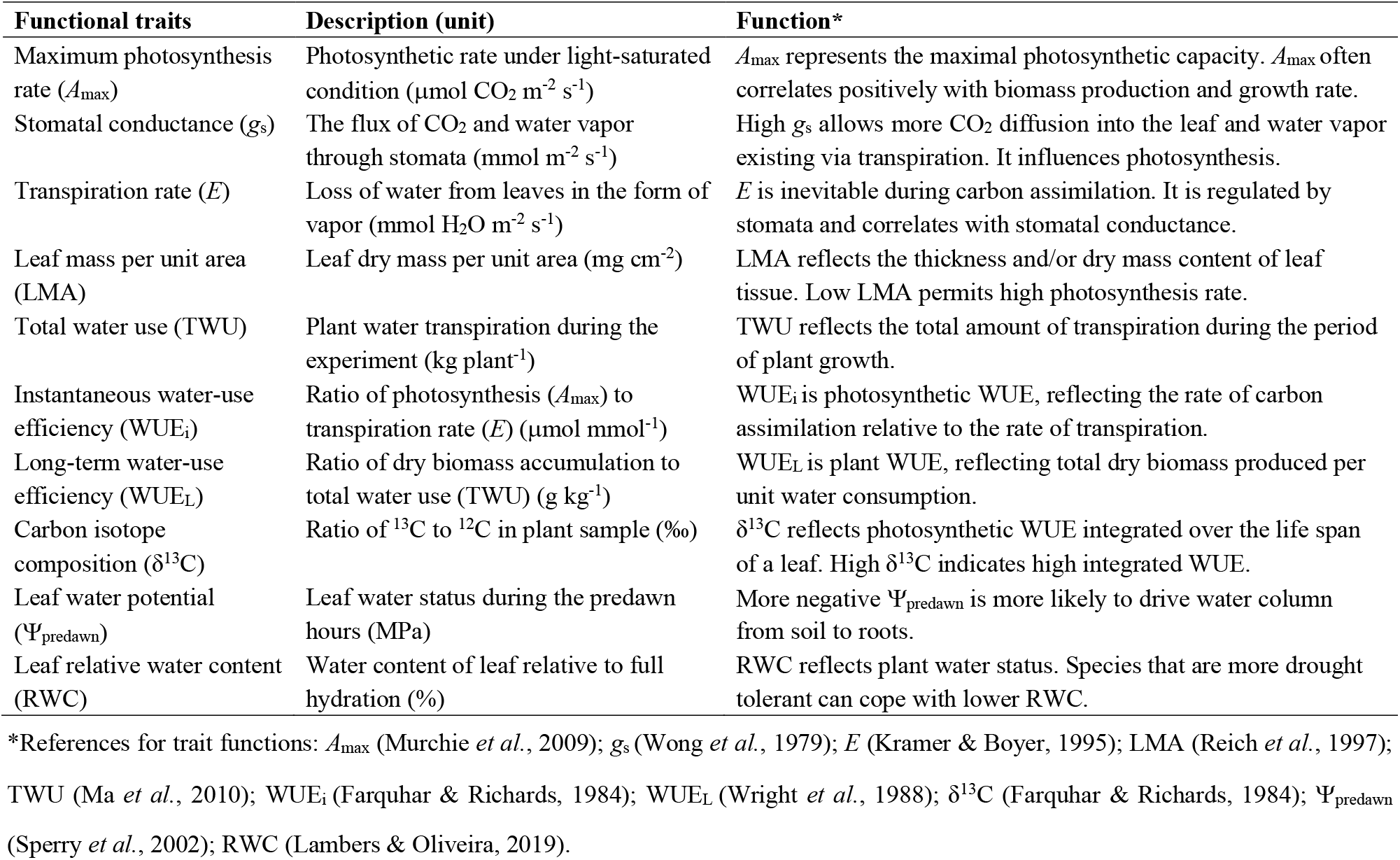
Key variables of functional traits

Functional traits were measured primarily at the end of the experiment during July 20–30, 2016. Gas exchange (maximum photosynthetic rate, *A*_max_; stomatal conductance, *g*_s_; transpiration rate, *E*) was measured for a random subset of 13 pots per ploidy level per treatment using a LI-COR LI-6400 portable photosynthesis system (Lincoln, NE, USA). Measurements were taken on a fully expanded green leaf from one plant each pot between 09:30 and 12:00 h at a saturating irradiance (1200 μmol m^−2^ s^−1^). The cuvette CO2 concentration was maintained at 400 μmol mol^−1^. Instantaneous water use efficiency (WUE_i_) was calculated as *A*_max_/*E*.

From the same subset of pots for gas exchange measurements, 10 pots per ploidy level per treatment were selected for leaf mass per unit area (LMA) and relative water content (RWC) measurements. Specifically, the largest, fully expanded green leaf was taken from one plant each pot and weighted immediately for fresh mass. Leaf area was scanned and estimated using ImageJ v1.45 (Schneider *et al*., 2012). The same leaf was then submerged in distilled water overnight to obtain the saturated mass, and dried at 80°C for 48 h for dry mass. LMA was calculated as leaf dry mass/leaf area, and RWC was calculated as (fresh mass – dry mass)/(saturated mass – dry mass) ×100. In addition, we measured predawn leaf water potential (Ψ_predawn_) using a WP4C Dewpoint Potential Meter (METER Group, Inc., Pullman, WA, USA) and carbon isotope composition (δ^13^C) in eight pots per ploidy level per treatment. δ^13^C was analyzed at the Institute of Tibetan Plateau Research.

Plant total water use (TWU) and long-term water use efficiency (WUE_L_) were measured in a different subset of 10 pots per ploidy level per treatment, to avoid the influence of destructive leaf collection described above. As the experimental and ‘empty’ pots were weighted every two days to record water loss (for maintaining specific water content), total water use each pot was determined as the difference between accumulated water loss each experimental pot (transpiration and evaporation) and the average accumulated water loss per empty pot (evaporation). Plant TWU was half of total water use each pot, due to two plants growing together. To determine WUE_L_, plants each pot were harvested for estimating dry biomass (including both belowground and aboveground) per plant. WUE_L_ was calculated as the ratio between plant dry biomass and TWU.

### Relative interaction intensity (RII)

The competitive interaction between plants was quantified using RII (Armas *et al*., 2004): RII = (B_w_ – B_o_)/(B_w_ + B_o_), where B_w_ and B_o_ are the plant biomass with and without competitors, respectively. RII ranges between −1 and 1, with RII < 0 indicting competition. In this study, B_w_ was the average plant biomass of the 2*x* or 4*x* each pot. Due to the lack of a single 2*x* or 4*x* plant growing alone here, B_o_ was calculated as the average plant biomass (with one intraploidy competitor present) across replicated pots for the 2*x* and 4*x* separately under each water stress treatment. As a result, our estimate of RII was conservative.

### Statistical analyses

To test for competitive asymmetry between ploidy levels (Fig. 1), we performed general linear mixed models (LMMs) with RII as the response variable using the package lme4 (Bates *et al*., 2015) in R v3.6.2 (R Core Team, 2019). The LMMs were conducted separately for the low and high water stress treatments. The predictors included ploidy (2*x* or 4*x*), intraploidy density (2, 3, 4 plants per pot) and interploidy density (0, 2, 3, 4), as well as the two-way interactions (intraploidy density – ploidy, interploidy density – ploidy). The random effects included pot positions (column and row) in the competition experiment to account for microenvironment variation and the fact that the 2*x* and 4*x* plants grew together in the same pots (Fig. S1). We did not consider source populations in the random effects, because seeds from different populations were mixed at random and sowed for the competition experiment. Statistical significance (type III sums of squares) and least-squares means (LS means) of predictors were assessed using packages lmerTest (Kuznetsova *et al*., 2017) and emmeans (Lenth, 2019). The slopes of interploidy density and intraploidy density in the LMMs represented respective competitive effects (i.e. α_4*x*,2*x*_ and α_2*x*,2*x*_ for the 2*x*; α_2*x*,4*x*_ and α_4*x*,4*x*_ for the 4*x*). The significance of interploidy density – ploidy interaction indicates competitive asymmetry between the 2*x* and 4*x*. To further test whether competitive asymmetry is maintained or even enhanced under stress (Fig. 1b), interploidy competitive effect (α_2*x*,4*x*_ or α_4*x*,2*x*_) was compared between the low and high water stress treatments using the *t* statistic for the 2*x* and 4*x* separately. Likewise, post hoc *t* tests were used to compare intraploidy vs. interploidy competitive effects (Fig. 1c).

To test whether the 4*x* is competitively dominant with higher biomass consistently across water stress conditions, we performed a LMM with log-transformed aboveground dry biomass as the response variable. The predictors included ploidy, stress, intraploidy density and interploidy density, as well as the two-way interactions (ploidy – stress, intraploidy density – ploidy, interploidy density – ploidy, intraploidy density – stress, interploidy density – stress) and three–way interactions (intraploidy density −ploidy – stress, interploidy density – ploidy – stress). The random effects included the column and row of pots in the competition experiment.

To evaluate how the 4*x* and 2*x* differ in functional traits, we conducted LMMs for individual traits separately. In all the LMMs, response variables were power transformed if necessary to improve normality, with the optimal power parameter determined using the Box–Cox method in the package car (Fox & Weisberg, 2011). The predictors included ploidy and stress and their interaction. For the random effects, we used the row or column of pots in the functional traits experiment, whichever explained more variation in the LMMs, but not both due to difficulties in model convergence. We did not include source populations in the random effects, because source populations explained less variation than pot positions, and often influenced model convergence.

## Results

### Competitive asymmetry at both low and high water stress

Consistent with the competitive asymmetry hypothesis (Fig. 1a,b), the competitive effect of 4*x* on 2*x* (α_2*x*,4*x*_; Fig. 2a, blue solid line) was stronger than that of 2*x* on 4*x* (α_4*x*,2*x*_; Fig. 2a, red solid line; LMM, interploidy density – ploidy interaction, *F* = 12.4, *P* = 0.0007). Specifically, the competitive effect was 1.67-fold higher for the 4*x* relative to the 2*x* under low water stress (LS mean ± SE, α_2*x*,4*x*_ = −0.15 ± 0.01, α_4*x*,2*x*_ = −0.09 ± 0.01; Fig. 2b). Under high water stress, the competitive response (RII) weakened by moving upwards (Fig. 2a). This trend followed the expectation of the stress gradient hypothesis (Fig. 1a); yet, this reduction in RII was not statistically significant for both the 4*x* and 2*x* (LMM, stress main effect, *F* = 0.15, *P* = 0.70; stress – ploidy interaction, *F* = 0.07, *P* = 0.80). Likewise, water stress did not significantly reduce the competitive effect (stress low vs. high, LS mean contrast, α_2*x*,4*x*_ = −0.15 ± 0.01 vs. −0.14 ± 0.01, *t* = 0.11, df = 103, *P* = 0.46; α_4*x*,2*x*_ = −0.09 ± 0.01 vs. −0.06 ± 0.02, *t* = 1.37, df = 107, *P* = 0.09; Fig. 2b). As a result, the competitive asymmetry of 4*x* vs. 2*x* was maintained under high water stress (*F* = 20.8, *P* < 0.001; Fig. 2a,b).

**Fig. 2.**
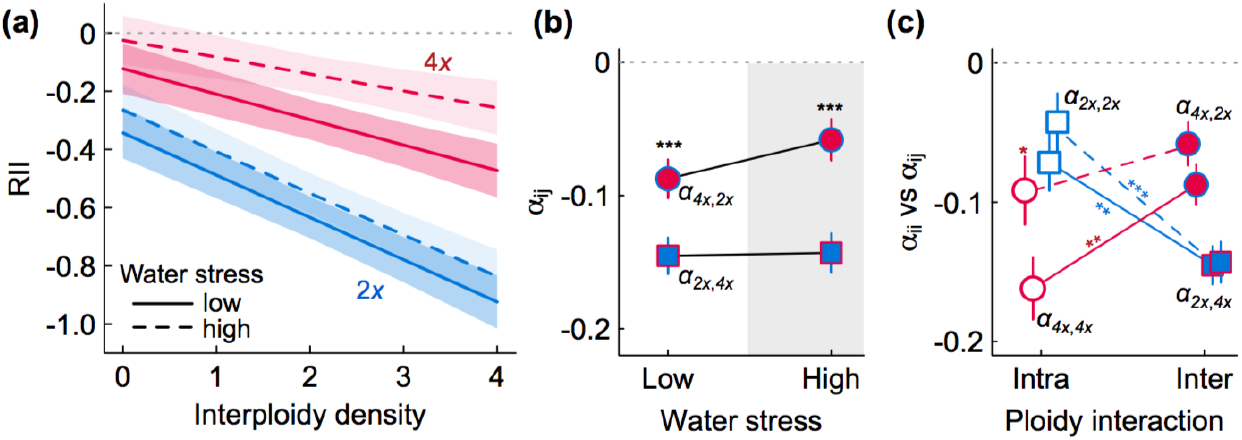
Experimental evidence for competitive asymmetry between polyploids and diploids. (a) Compeition experienced by the 4*x* and 2*x* (measured using the relative interaction index, RII) intensified as their interploidy competitors increased in density, as revealed by general linear mixed models (LMMs). The interploidy competitive effect (slope, with shaded 95% confidence intervals) was stronger on the 2*x* than the 4*x* under both low (LMM, *F* = 12.4, *P* = 0.0007, solid lines) and high water stress (*F* = 20.8, *P* = 0.00002, dashed lines). (b) The least-squares mean (LS mean) ± 1 SE of interploidy competitive effect (α_*ij*_, where *i* and *j* refer to the 2*x* or 4*x*, respectively) are plotted. (c) The LS mean ± 1 SE of intraploidy (α_*ii*_) vs. interploidy competitive effect (α_*ij*_ are plotted. Contrasts of LS means were based on post-hoc *t* tests, with significance levels labeled along the connecting lines or above symbols (for contrasts between vertical symbols): \*\*\**P* < 0.001; \*\**P* < 0.01; \**P* < 0.05.

Consistent with the expectation that intraploidy effect is stronger than interploidy effect in polyploids (Fig. 1c), the magnitude of α_4*x*,4*x*_ (LS mean ± SE, −0.16 ± 0.02) was nearly two-fold higher than α_4*x*,2*x*_ (−0.09 ± 0.01, *t* = 2.79, df = 106, *P* = 0.003; Fig. 2c) under low water stress. But this pattern was weakened under high water stress (α_4*x*,4*x*_ = −0.09 ± 0.02, α_4*x*,2*x*_ = −0.06 ± 0.02, *t* = 1.15, df = 108, *P* = 0.13), due to significantly reduced intraploidy competitive effect in the 4*x* under water stress (α_4*x*,4*x*_, LS mean contrast, *t* = 2.12, df = 107, *P* = 0.018), whereas the reduction in interploidy effect of 2*x* on 4*x* was not significant (α_4*x*,2*x*_, *t* = 1.37, df = 107, *P* = 0.09; Fig. 2c). For diploids, consistent with the expectation that intraploidy effect is weaker than interploidy effect (Fig. 1c), the magnitude of α_2*x*,2*x*_ (−0.07 ± 0.02) was half of α_2*x*,4*x*_ (−0.15 ± 0.01, *t* = 3.0, df = 100, *P* = 0.002) under low water stress (Fig. 2c), and this pattern held under high water stress (α_2*x*,2*x*_ = −0.04 ± 0.02, α_2*x*,4*x*_ = −0.14 ± 0.01, *t* = 3.96, df = 106, *P* < 0.001).

In addition to competitive asymmetry (Fig. 2), the competitive dominance of the 4*x* was also reflected by overall higher aboveground biomass under both low (*χ*^2^ = 223, df = 1, *P*< 0.001) and high water stress (*χ*^2^ = 262, df = 1, *P* < 0.001; Fig. 3).

**Fig. 3.**
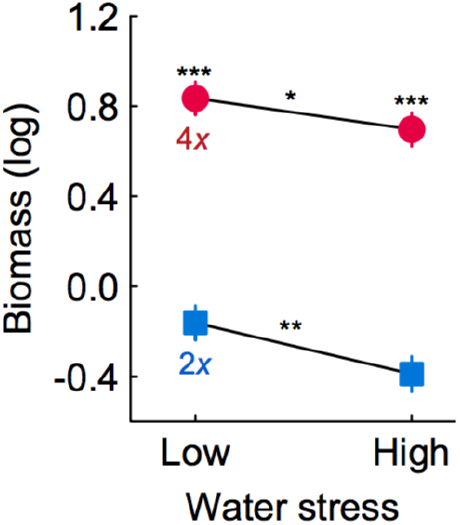
Polyploids are competitively dominant under different water stress conditions in the competition experiment. The aboveground dry biomass was log transformed in the general linear mixed model (LMM). The least-squares mean (LS mean) ± 1 SE of the 2*x* (blue squares) and 4*x* (red circles) are plotted. Significant contrasts of LS means are denoted along the connecting lines or above symbols (for contrasts between vertical symbols): \*\*\**P* < 0.001; \*\**P* < 0.01; \**P* < 0.05.

### Functional traits explaining competitive dominance of polyploids

Under low water stress, the 4*x* showed more resource-acquisitive traits relative to the 2*x* (Fig. 4a-e), including higher maximum photosynthesis rate (*A*_max_, LS mean ± SE, 4*x* = 12.1 ± 0.5, 2*x* = 10.3 ± 0.5, *χ*^2^ = 18.2, df = 1, *P* < 0.001), stomatal conductance (*g*_s_, 4*x* = 0.40 ± 0.02, 2*x* = 0.26 ± 0.02, *χ*^2^ = 63.6, df = 1, *P* < 0.001), transpiration rate (*E*, 4*x* = 3.7 ± 0.2, 2*x* = 3.3 ± 0.2, *χ*^2^ = 5.0, df = 1, *P* = 0.025) and total water use (TWU, 4*x* = 1.4 ± 0.1, 2*x* = 0.7 ± 0.1, *χ*^2^ = 36.0, df = 1, *P* < 0.001), as well as lower leaf mass per area (LMA, 4*x* = 2.2 ± 0.1, 2*x* = 2.6 ± 0.1, *χ*^2^ = 11.0, df = 1, *P* = 0.0009). Meanwhile, the 4*x* and 2*x* were similar in water use efficiency (WUE_i_, *χ*^2^ = 0.002, df = 1, *P* = 0.96; WUE_L_, *χ*^2^ = 1.02, df = 1, *P* = 0.31; δ^13^C, *χ*^2^ = 1.12, df = 1, *P* = 0.29; Fig. 4f-h) and leaf water potential (Ψ_predawn_, *χ*^2^ = 0.53, df = 1, *P* = 0.47; Fig. 4i) and relative water content (RWC; *χ*^2^ = 0.09, df = 1, *P* = 0.76; Fig. 4j).

**Fig. 4.**
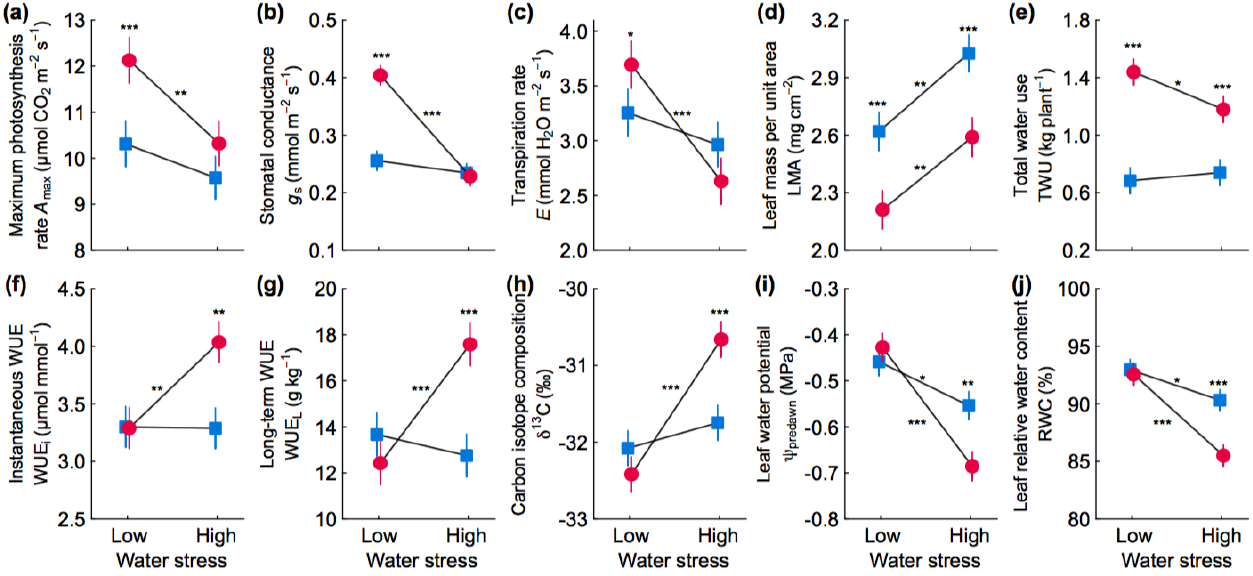
Polyploids are more resource aquisitive under low water stress and more drought tolerant under high water stress than diploids. The least-squares mean (LS mean) ± 1 SE of individual functional traits (Table 1) are plotted for the 2*x* (blue squres) and 4*x* (red circles), estimated from general linear mixed models (LMMs). WUE, water use efficiency. Significant contrasts of LS means are denoted along the connecting lines or above symbols (for contrasts between vertical symbols): \*\*\**P* < 0.001; \*\**P* < 0.01; \**P* < 0.05.

In contrast, under high water stress, the 4*x* showed more drought tolerant traits relative to the 2*x* (Fig. 4f-j), including higher water use efficiency (WUE_i_, 4*x* = 4.0 ± 0.2, 2*x* = 3.3 ± 0.2, *χ*^2^ = 9.3, df = 1, *P* = 0.002; WUE_L_, 4*x* = 17.6 ± 0.9, 2*x* = 12.8 ± 0.9, *χ*^2^ = 15.4, df = 1, *P* = 0.0001; δ^13^C, 4*x* = −30.7 ± 0.2, 2*x* = −31.7 ± 0.2, *χ*^2^ = 11.4, df = 1, *P* = 0.0007), and lower Ψ_predawn_ (4*x* = −0.69 ± 0.03, 2*x* = −0.55 ± 0.03, *χ*^2^ = 9.4, df = 1, *P* = 0.002) and RWC (4*x* = 86% ± 1%, 2*x* = 90% ± 1%, *χ*^2^ = 14.3, df = 1, *P* = 0.0002), while keeping similar levels of gas exchange as the 2*x* (*A*_max_, *χ*^2^ = 2.9, df = 1, *P* = 0.09; *g*_s_, *χ*^2^ = 0.1, df = 1, *P* = 0.75; *E*, *χ*^2^ = 2.8, df = 1, *P* = 0.10).

## Discussion

By quantifying the interaction intensity of interploidy and intraploidy competitions, our results provide strong evidence for the competitive asymmetry between polyploid and diploid plants in *C. indicum*, with the tetraploid that is geographically more widespread being competitively dominant to the diploid. As a stronger competitor, the tetraploid was also found to be limited more by individuals of its own than individuals of the diploid, and the predicted reverse pattern was detected in the diploid. Such competitive asymmetry was not only maintained under increased water stress, but also translated into higher fitness of the tetraploid than the diploid consistently across water stress conditions, demonstrating a ‘jack-and-master’ fitness strategy (see also Wei *et al*., 2019) of the tetraploid in *C. indicum*. Our results further reveal that functional traits associated with fast growth and efficient water use likely underlie the competitive dominance of tetraploid *C. indicum*.

### Polyploids are competitively dominant relative to diploids

Our results of stronger interploidy competitive effect of the tetraploid than the diploid support the hypothesis that competition between polyploid and diploid individuals is asymmetric in *C. indicum*. The competitive abilities of polyploids and diploids have previously been evaluated by two different methods: ‘direct’ assessment where competing polyploids and diploids grow together (polyploid–diploid competition) (e.g. ours here; Maceira *et al*., 1993; Thompson *et al*., 2015; Rey *et al*., 2017) and ‘indirect’ assessment where polyploids and diploids do not compete with each other but instead each competes with other plant species (polyploid–other species vs. diploid–other species competition) (Čertner *et al*., 2019). The direct assessment can inform the strength of competitive effect that one ploidy may exert on another when encountering each other (Maceira *et al*., 1993), whereas the indirect assessment considers the possibility that co-occurring polyploids and diploids may have other more immediate neighbors (Čertner *et al*., 2019). Similar to our findings, previous studies using direct assessments have revealed stronger competitive effects of polyploids than diploids in grass *Dactylis glomerata* (Maceira *et al*., 1993), forb *Centaurea stoebe* (Collins *et al*., 2011) under low stress conditions. Likewise, indirect assessments have revealed lower fitness loss of polyploids than diploids when competing with other plant species in herbaceous *Lolium perenne* (Sugiyama, 1998), *Knautia serpentinicola* (Čertner *et al*., 2019), and *Solidago canadensis* (Cheng *et al*., 2020). These previous studies and ours provide empirical evidence for the competitive dominance of polyploids across different plant systems.

Also consistent with our predictions, the competitively dominant tetraploid *C. indicum* experienced stronger intraploidy than interploidy competition especially under low water stress in this study. This pattern was seen previously in tetraploid *D. glomerata* (Maceira *et al*., 1993) where intraploidy competitive coefficient was stronger than interploidy (α_4*x*,4*x*_ > α_4*x*,2*x*_), and the predicted reverse pattern (α_2*x*,2*x*_ < α_2*x*,4*x*_) was also detected in its diploid. In tetraploid *Brachypodium hybridum* (Rey *et al*., 2017), α_4*x*,4*x*_ > α_4*x*,2*x*_ was also detected; but the prediction of α_2*x*,2*x*_ < α_2*x*,4*x*_ in its diploid (*B. distachyon*) was environment dependent and was only supported in drier environments. Different from these findings, in *Chamerion angustifolium* (Thompson *et al*., 2015), similar levels of intraploidy and interploidy competition coefficients were observed in both the tetraploid and diploid plants. These differences in empirical support for intra- vs. interploidy competition indicate the complexity of species interaction and its context dependence, which can vary with other factors, for instance, experimental environments and pre-existing adaptation of polyploid and diploid populations to the experimental environments. Thus, additional research is needed for a more complete understanding of species interactions between and within ploidy levels across diverse polyploid–diploid systems and environments.

### Competitive asymmetry is maintained under high water stress

Our results follow the trend predicted by SGH that competitive interactions weaken under stress (Fig. 2), albeit not yet statistically significant. While the tetraploid and diploid plants were negatively affected by water stress in terms of aboveground biomass (Fig. 3), the lack of significant difference in competitive response (RII) and effect (competitive coefficients) between low and high water stress conditions likely have several reasons. First, plants may still experience strong competition despite that they were smaller in size under high water stress, given the limited space and resource of fixed-size pots in this study. Second, the competitive response and effect measured in this study focused on one important fitness component, vegetative growth, in perennial *C. indicum*. While vegetative robustness in perennial plants likely positively affects sexual reproduction as well (Fujita *et al*., 2014; Wei *et al*., 2019; Cheplick, 2020), it remains to be evaluated how competitive response and effect based on other fitness components (e.g. sexual reproduction) follow the predictions of SGH. For instance, in line with SGH, the interploidy competitive effects of annual grass diploid *B. distachyon* and tetraploid *B. hybridum* measured based on seed production decreased in drier conditions (Rey *et al*., 2017). Third, the level of experimental stress and how plants cope with the imposed stress can also influence the outcome of species interactions. As *C. indicum* especially the tetraploid populations often occur in dry habitats (Li *et al*., 2014), they likely have evolved strategies to cope with severe stress in the wild, which might be more severe than a constant soil moisture of 20% field capacity here (Fig. 4). Thus, as factors including neighbor density, resources availability and stress tolerance can influence the strength of species interactions along stress gradients (Donovan & Richards, 2000; Liancourt *et al*., 2005; Maestre *et al*., 2009; Zhang & Tielböerger, 2019, 2020), it is important to consider these factors when evaluating SGH in plants including the polyploid–diploid systems.

Given the limited influence of stress on interploidy competitive effect here, the competitive asymmetry between tetraploid and diploid *C. indicum* was maintained under high water stress. While competitive asymmetry was found to vary with environments between diploid *B. distachyon* and tetraploid *B. hybridum* (Rey *et al*., 2017), the maintenance of competitive asymmetry in *C. indicum* under both low and high water stress conditions translated into higher fitness of the tetraploid than the diploid. Such ‘jack-and-master’ fitness strategy (Wei *et al*., 2019) of tetraploid *C. indicum* may underlie their widespread distribution across heterogeneous environments relative to diploid *C. indicum* (Li *et al*., 2014). As more intense and frequent drought events are happening under global climate change (Ploughe *et al*., 2019), interactions within and between diploid and tetraploid *C. indicum* populations might respond differently, which can influence their future distributions.

### Functional traits explain the competitive dominance of tetraploid *C. indicum*

Tetraploid *C. indicum* expressed traits that can confer faster growth under low water stress and more efficient water use under high water stress relative to the diploid, which may underlie the competitive dominance of the tetraploid. Under low water stress, the acquisitive strategy of the tetraploid was manifested by higher gas exchange and higher total water use. Interestingly, different from many other polyploid–diploid systems (e.g. Li *et al*., 2009; Greer *et al*., 2018; Wei *et al*., 2019), tetraploid *C. indicum* had lower LMA (higher SLA), which may contribute to greater photosynthetic capacity and higher biomass accumulation (Reich *et al*., 1997; Westoby *et al*., 2002). Under high water stress, the tetraploid exhibited higher water use efficiency, which can benefit the tetraploid in conserving water for growth under water deficiency (Manzaneda *et al*., 2015; Greer *et al*., 2018). In addition, the tetraploid was able to maintain gas exchange at a similar level as the diploid but at a lower level of leaf water potential (Ψ_predawn_) and leaf relative water content (RWC), indicating stronger drought tolerance of the tetraploid, similar to what has been observed in polyploid *Betula papyrifera* (Li *et al*., 1996). Different from drought tolerance in tetraploid *C. indicum*, drought avoidance was found in polyploid *C. angustifolium* (Maherali *et al*., 2009), *Lonicera japonica* (Li *et al*., 2009) and *Citrus sinensis* (Oliveira *et al*., 2017), where the polyploids exhibited higher Ψ_predawn_ and RWC under drought. Overall, functional traits associated with fast growth and efficient water use likely underlie the competitive dominance of tetraploid *C. indicum* and its more widespread distribution.

To conclude, our study provides a quantitative demonstration of the competitive asymmetry between polyploids and diploids consistently across stress conditions. This offers an important mechanistic insight into the ecological advantage of polyploids (Wei *et al*., 2019; Van de Peer *et al*., 2021) and the underlying functional traits. Our study awaits similar investigations on other plant lineages to generalize the species interactions between polyploid and diploid plants across heterogeneous environments, and together shed light on population dynamics and species distribution in the face of environmental change.

## Acknowledgements

This work was supported by the Second Tibetan Plateau Scientific Expedition and Research (STEP) program (2019QZKK0502), the National Natural Science Foundation of China (31800334 to WG, 31760114 to SJY) and the Fundamental Research Projects from Science and Technology Department of Yunnan Province (2019FD004 to WG, 202101AT070180 to SJY). We acknowledge the Kunming Institute of Botany glasshouse for logistic support.

## Author contributions

WG, YWD and YPY designed the research, and NW conceived the conceptual development of the manuscript. WG conducted the glasshouse experiments. NW analyzed the data. WG and NW wrote the manuscript, and all authors contributed to revisions.

## Supporting Information

**Table S1.** Sampling information of diploid and tetraploid *Chrysanthemum indicum*.

| Site name | SiteID | PopID | Latitude<br>(°N) | Longitude<br>(°E) | Altitude<br>(m) | Ploidy | N<br>* |
| --- | --- | --- | --- | --- | --- | --- | --- |
| Songlangshan | Pop.2x.1 | Pop.2x.1.1 | 31.75026 | 110.67218 | 1054 | 2x | 5 |
| Panshuihe | Pop.2x.2 | Pop.2x.2.1 | 31.75558 | 110.60596 | 1163 | 2x | 3 |
| Songbaicun | Pop.2x.3 | Pop.2x.3.1 | 31.75397 | 110.65366 | 970 | 2x | 3 |
| Songbaicun | Pop.2x.3 | Pop.2x.3.2 | 31.75437 | 110.65356 | 1003 | 2x | 3 |
| Gumiaoya | Pop.4x.1 | Pop.4x.1.1 | 31.74358 | 110.60515 | 1265 | 4x | 6 |
| Baihuaping | Pop.4x.2 | Pop.4x.2.1 | 31.75346 | 110.65283 | 957 | 4x | 6 |
| Muyuzhen | Pop.4x.3 | Pop.4x.3.1 | 31.47910 | 110.38218 | 1241 | 4x | 8 |
| Muyuzhen | Pop.4x.3 | Pop.4x.3.2 | 31.48138 | 110.38182 | 1241 | 4x | 8 |
| Qingshancun | Pop.4x.4 | Pop.4x.4.1 | 31.49070 | 110.35798 | 1824 | 4x | 10 |
| Qingshancun | Pop.4x.4 | Pop.4x.4.2 | 31.49943 | 110.36758 | 1576 | 4x | 8 |

\*The number of maternal plants from which seeds were collected varied with population sizes.

**Fig. S1.**
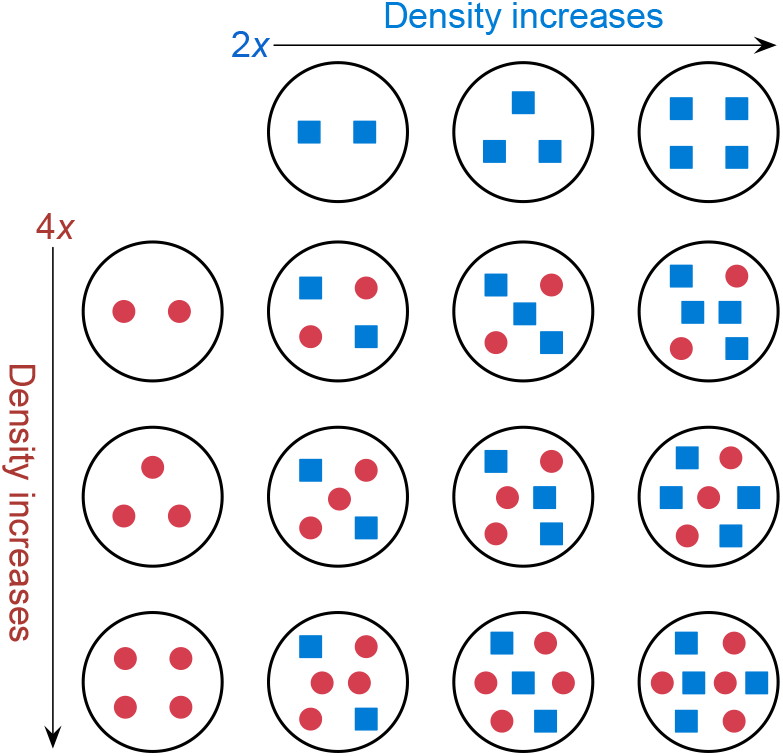
Competition experiment design. The basic unit of the competition experiment consists of 15 combinations that vary in the density of diploid (2x; blue squares) and tetraploid (4x; red circles) plants. This basic unit of the competition experiment was replicated at both low water stress and high water stress conditions, each with 5 replicates.

